# Driving Native-like Zonal Enthesis Formation in Engineered Ligaments Using Mechanical Boundary Conditions and β-Tricalcium Phosphate

**DOI:** 10.1101/2021.07.24.453656

**Authors:** M. Ethan Brown, Jennifer L. Puetzer

## Abstract

Fibrocartilaginous entheses are structurally complex tissues that translate load from elastic ligaments to stiff bone via complex zonal organization with gradients in organization, mineralization, and cell phenotype. Currently, these gradients, necessary for long-term mechanical function, are not recreated in soft tissue-to-bone healing or engineered replacements, leading to high failure rates. Previously, we developed a culture system which guides ligament fibroblasts to develop aligned native-sized collagen fibers using high density collagen gels and mechanical boundary conditions. These constructs hold great promise as ligament replacements, however functional ligament-to-bone attachments, or entheses, are required for long-term function *in vivo*. The objective of this study was to investigate the effect of compressive mechanical boundary conditions and the addition of beta tricalcium phosphate (βTCP), a known osteoconductive agent, on the development of zonal ligament-to-bone entheses. We found that compressive boundary clamps, that restrict cellular contraction and produce a zonal tensile-compressive environment, guide ligament fibroblasts to produce 3 unique zones of collagen organization, and zonal accumulation of glycosaminoglycans (GAGs), type II and type X collagen by 6 weeks of culture, ultimately resulting in similar organization and composition as immature bovine entheses. Further, βTCP under the clamp enhanced the maturation of these entheses, leading to increased GAG accumulation, sheet-like mineralization, and significantly improved tensile moduli, suggesting the initiation of endochondral ossification. This culture system produced some of the most organized entheses to date, closely mirroring early postnatal enthesis development, and provides an *in vitro* platform to better understand the cues that drive enthesis maturation *in vivo*.

## 1. Introduction

Ligaments are anchored into bone via insertion sites known as entheses. Entheses are structurally complex tissues 100 µm to 1 mm wide that act to translate load from elastic ligaments to stiff bone via a compliant fibrocartilage region [1–4]. Entheses withstand and distribute these complex loads due to a zonal organization with spatial gradients in organization, composition, mineralization, and cell type (**Figure 1**). Specifically, this tissue is marked by aligned collagen fibers in the ligament to more randomly oriented fibers with increasing mineral and glycosaminoglycan (GAG) content as you move from fibrocartilage to bone. Similar to these gradients in matrix composition there are gradients in cell type, consisting of fibroblasts, fibrochondrocytes, hypertrophic chondrocytes, osteoblasts, osteoclasts, and osteocytes from the ligament to the bone, respectively [1–4].

**Figure 1:**
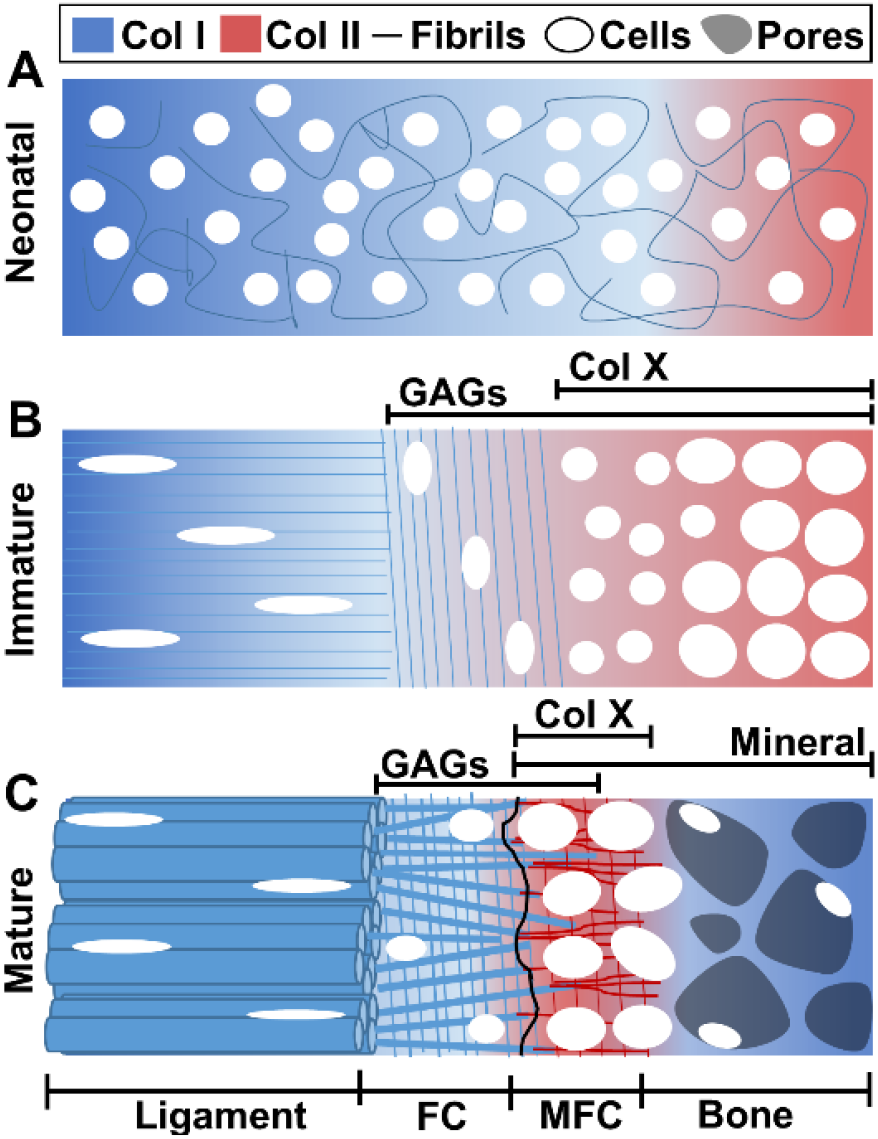
Depiction of enthesis development from A) neonatal to B) immature to C) mature tissue, resulting in continuous gradients of organization, composition, and cell phenotypes across ligament, fibrocartilage (FC), mineralized fibrocartilage (MFC), and bone in mature tissue

Currently, these gradients of the enthesis, necessary for long-term mechanical function, are not recreated in soft tissue-to-bone healing or engineered replacements [2–4]. Each year there are approximately 225,000 anterior cruciate ligament (ACL) injuries in the US alone [5], of which about 135,000 ACL reconstructions are performed with an estimated economic burden of $5.1 billion [3,6,7]. Further, 10-20% of ACL graft repairs fail within 5 years [8–10], primarily at the enthesis, due to a lack of regeneration [3,11]. It remains a challenge to form the physiological gradients of the enthesis necessary for mechanical function [1–3]. Engineered tissues are a promising solution to repair ACLs, however, attempts at replacing the ACL have historically focused on the ligament proper [12–14] and lack functional entheses for proper anchoring to bone [15–20]. More recent efforts at ligament-to-bone, or ligament-interface-bone replacements have primarily focused on top-down scaffold design using aligned synthetic or natural fibers and induced spatial mineralization [21–27]. These techniques attempt to replicate mature ligament and bone organization; however, they may actually impede development of the physiological gradient of the enthesis since aligned fibers and mineralization are not present until later in development.

Recapitulating development (**Figure 1**, based on [28–36]), in a bottom-up approach, is a promising means to produce enthesis gradients. Initially, the enthesis is primarily composed of homogenously distributed type I collagen fibrils (**Figure 1A**) [28]. Postnatally, a fibrocartilage interface develops with evolving gradients in cells, collagen types I, II and X, proteoglycans, and mineralization. Notably, the collagen organization in the enthesis transitions from unorganized at birth to primarily perpendicular to the ligament during postnatal development (**Figure 1B**) [25,32,35]. Finally, as the enthesis matures, collagen shifts from being perpendicular to the ligament to a more diffuse orientation parallel with the ligament (**Figure 1C)**, with oblique insertions into the bone [25,35,36]. This final collagen organization is thought to be critical to reducing stress concentrations and functional translation of load [30,31,33,34,37].

The enthesis-to-bone transition develops primarily by endochondral ossification [2,4,28,32]. The bone region is initially cartilage-like, composed of chondrocytes, type II collagen, and proteoglycans, such as aggrecan (**Figure 1A)** [36]. With development these chondrocytes become hypertrophic, eventually forming bone primarily composed of type I collagen and mineral (**Figure 1B**). During postnatal development, type II collagen, type X collagen, and GAG accumulation all shift from the bone region to the fibrocartilage region of the enthesis (**Figure 1C**) [1,3,4,33]. Mineralization is not present until late postnatal development and is thought to form a composite with the collagen fibers, increasing the toughness of the attachment [29,38–40].

Recently, we developed a novel culture device (**Figure 2A**), which uses compressive boundary restraints (clamps) to guide ligament fibroblasts in high density collagen gels to form native sized and aligned 30 µm diameter collagen fibers which group together to form ∼200 µm diameter fiber bundles by 6 weeks of culture [41]. These constructs, formed through a bottom-up approach, have some of the largest and most organized hierarchical collagen fibers produced to date *in vitro*, holding great promise as engineered ligament replacements. However, driving organized enthesis development is critical for proper function *in vivo*.

**Figure 2:**
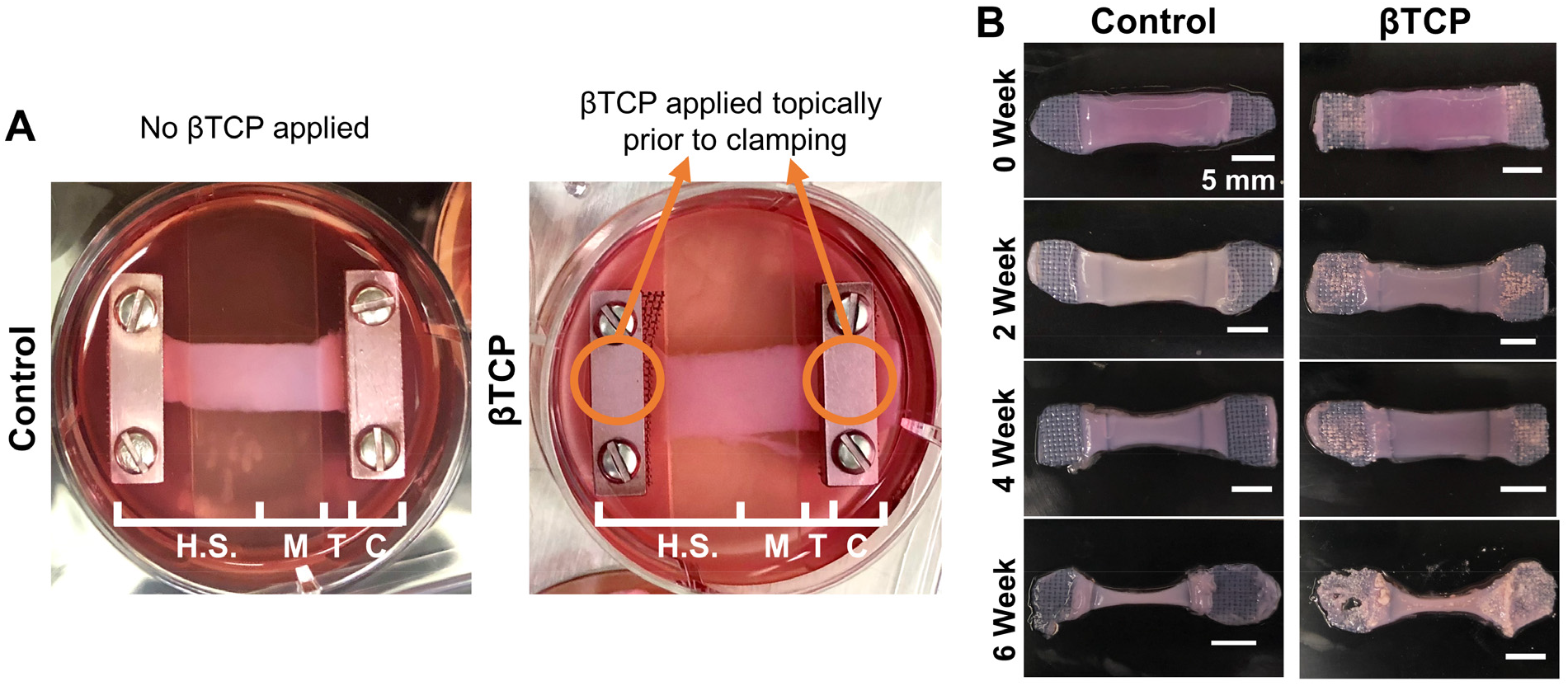
A) Cell-seeded high density collagen gels in clamping device at 0 weeks depicting control group and βTCP group. Depiction of half-sample (H.S.) and Middle (M), Transition (T), Clamped (C) regions obtained for analysis; B) Photographs of constructs at each time point depicting similar gross morphological changes throughout culture. Scale bar = 5 mm.

A major feature of this culture system is the formation of a unique interface between the aligned midsection and the compressive clamps [41]. Specifically, this culture device has a 1 mm step down to the clamped boundary, producing a compressive environment under the clamps and a tensile-compressive environment between the clamps and the midsection. Traditionally, tensile loads are often fibrogenic, increasing type I collagen synthesis and tensile properties, while compressive loads are chondrogenic, increasing GAG and type II collagen synthesis [42,43]. Further, dual tensile-compressive loads have been shown to drive fibrocartilage-like tissue with increased GAG, type I collagen, and type II collagen synthesis in engineered menisci [44,45]. Therefore, we hypothesize the tensile-compressive environment of our culture device will drive fibrocartilage enthesis-like tissue that shifts to cartilage-like tissue under the compressive clamp, producing a ligament-to-bone transition similar to neonatal enthesis tissues.

In an effort to further develop the region under the clamp to a more bone-like tissue, we also investigated the addition of beta tricalcium phosphate (βTCP) under the clamp. βTCP is known to be bioresorbable and osteoconductive [46,47], and further has been demonstrated to release a greater amount of calcium ions *in vitro* than hydroxyapatite [48]. The addition of calcium phosphate mineral has been shown to enhance the interface between soft and hard tissues [25,26,49]. It is thought that complete incorporation of tissue engineered bone-ligament-bone only occurs when the mineral used in the engineered tissue is replaced with endogenous bone [25] indicating the use of bioresorbable calcium phosphates such as βTCP as a logical option over slower degrading hydroxyapatite [47,50]. The addition of βTCP under the boundary clamps (**Figure 2A**) will yield a contained, osteogenic environment for the cells throughout the duration of culture, yielding two unique culture conditions between the clamps and under the clamps.

The objective of this study was to investigate the effect of compressive mechanical boundary conditions and the addition of βTCP on the development of zonal ligamentous entheses, specifically evaluating zonal collagen organization, matrix composition, and mechanics. We hypothesize the tensile-compressive interface and compressive clamps will produce zonal fibrocartilage and cartilage-like tissue formation similar to early-postnatal enthesis, and the addition of βTCP will further stimulate the tissue under the clamp to be more bone-like, producing engineered entheses with zonal organization and matrix composition.

## 2. Methods

### 2.1 Calcium Phosphate Synthesis

Beta tricalcium phosphate (βTCP) was synthesized via a wet chemical precipitation reaction of calcium nitrate tetrahydrate and diammonium hydrogen phosphate as previously described [46]. The precipitate was filtered, dried, and calcined at 900°C for two hours. The final calcined βTCP was ball milled into a fine powder to permit for localized delivery. To confirm βTCP synthesis X-ray diffraction (XRD) and Fourier-transform infrared (FTIR) spectroscopy were performed to characterize the phase and functional groups, respectively. XRD of the synthesized βTCP revealed characteristic peaks, as compared to JCPDS card number 09-0169, and FTIR displayed known phosphate absorption bands (**Supplemental Figure 1**) [46].

### 2.2 Construct Fabrication

#### 2.2.1 Cell Isolation

Ligament fibroblasts were isolated as previously described [41,44,51,52]. Briefly, ACLs were aseptically isolated from immature bovine legs (0-5 months old) obtained from a local slaughterhouse. The ligaments were diced and digested in 0.2% w/v collagenase (Worthington) overnight [53]. Cells were then filtered, washed, and counted. Isolated ligament fibroblasts were seeded at 2800 cells/cm^2^ and passaged 1-2 times to yield sufficient cell numbers. Two isolations were performed, with each isolation consisting of 3-4 donors combined into a single cell suspension for creation of constructs, which were then divided equally between control and βTCP constructs.

#### 2.2.2 Collagen Gel Fabrication and Culturing

Cell seeded high density collagen gels were fabricated as previously described [41,44,51,52]. In short, type I collagen was extracted from an equal amount of male and female Sprague Dawley rat tail tendons (BIOIVT) and reconstituted at 30 mg/mL in 0.1% acetic acid [41,54]. The 30 mg/mL collagen solution was mixed with a working solution of NaOH and PBS to initiate gelation and raise the pH and osmolarity to 7.0 and 300 mOsm, respectively. This collagen solution was immediately mixed with ligament fibroblasts to ensure cells were seeded throughout the collagen gel [41,44,52,54]. The cell-collagen solution was injected between glass sheets 1.5 mm apart and gelled at 37°C for 1 hour to obtain final constructs at 20 mg/mL collagen concentration with 5 million cells/mL. Rectangular constructs (30 x 8 mm) were cut from the final sheet gels using a die. Constructs were clamped on day 1 with βTCP constructs receiving 2-3 mg βTCP applied topically directly under each clamp, and controls receiving no βTCP. A piece of steel mesh was placed directly under the clamp to facilitate nutrient diffusion to the cells, as well as to prevent slippage of the constructs throughout culture [41]. The constructs were cultured in media consisting of Dulbecco’s modified Eagle media (DMEM), 10% fetal bovine serum, 1% antibiotic/antimycotic, 50 µg/mL ascorbic acid, and 0.8 mM L-proline, changed every 2-3 days. Constructs were cultured for up to 6 weeks with time points taken at 0, 2, 4, and 6 weeks. Zero week constructs were harvested 24 hours after clamping and application of βTCP to evaluate initial effects. A total of N = 7-11 constructs were harvested at each timepoint per condition, photographed, and sectioned into half-length samples or middle, transition, and clamped (M, T, and C) regions (**Figure 2A**) for analysis of zonal organization, composition, and mechanical properties.

### 2.3 Zonal Collagen Organization Analysis

Half-length sections, including middle, transition, and clamped regions, were fixed in 10% formalin and stored in 70% ethanol. Analysis was performed using scanning electron microscopy (SEM), confocal reflectance, and picrosirius red staining imaged with polarized light to evaluate hierarchical collagen organization at the fibril (<1 µm length scale), fiber (1-100 µm length scale), and fascicle (>100 µm length scale) length-scales, respectively, as previously reported [41]. A total of N = 4-11 constructs per time point and condition were analyzed via confocal reflectance. Following confocal analysis, a subset of 6 week constructs were imaged via SEM (N = 3) and picrosirius red (N = 4), resulting in a higher N (N = 10-11) for 6 week constructs compared to 0, 2, and 4 week constructs (N = 4-6). Engineered tissues were compared to native 4-5 months old bovine ACL-to-bone tissues, with the middle, transition, and clamped regions of engineered tissues compared to native ligament, fibrocartilaginous enthesis, and bone regions, respectively.

#### 2.3.1 Confocal Imaging Analysis

Confocal analysis was performed as previously described [41,44,51,52] with a Zeiss 710 inverted laser scanning microscope and LD C-Apochromat 40x/1.1 W Corr M27 objective. Briefly, a 488 nm laser was split between confocal reflectance and fluorescence. Collagen was visualized at 400-465 nm by collecting backscattered light through a 29 µm pinhole with pixel dwell time of 0.79 µs and cells were imaged via auto-fluorescence at 509-571 nm. Representative images were taken across the full length of each construct, with three to five images taken in each of the three zones for all samples. All images were then analyzed using FIJI (ImageJ, NIH) to assess collagen fiber alignment and dispersion. Briefly, the Despeckle 3 x 3 pixel median filter was applied to each image, followed by Fourier components analysis via the Directionality plugin. Collagen fiber direction data was binned every 2° from -90° to 90°, with 0° oriented horizontally across the image and 90° oriented vertically, to form a histogram of fiber direction counts. The histogram for each image was fit with a two-term Gaussian curve. Maximum collagen fiber alignment for each image is reported as the maximum of the histogram and dispersion is reported as the standard deviation of the gaussian fit. The absolute value was taken for any direction angles measuring between 0 and -90°. Images without any directionality resulted in flat histograms with very large standard deviations which were capped at a dispersion value of 90°. The resulting maximum angles of collagen fiber alignment per image were binned at 10° increments and are represented as percentage of images per bin from 0° to 90° (**Figure 3C**). The average dispersion of collagen fibers for each construct was determined by averaging all images for each region (n = 3-5 images per region per construct), with the overall dispersion consisting of the average of all constructs per timepoint (N = 4-11). Degree of collagen fiber dispersion is represented as a single wedge measured from 0° (**Figure 3C**). Overall averages for maximum angle of fiber alignment were calculated similarly and represented both by rose plots displaying percentages of images per 10° alignment (**Figure 3C**) and as bar graphs representing average maximum angle of fiber alignment (**Supplemental Figure 2**). Statistical analyses were performed for both maximum collagen fiber alignment and dispersion using overall averages, with statistical symbols presented in the 0° to 90° data displays (**Figure 3**) and bar graphs (**Supplemental Figure 2**).

**Figure 3:**
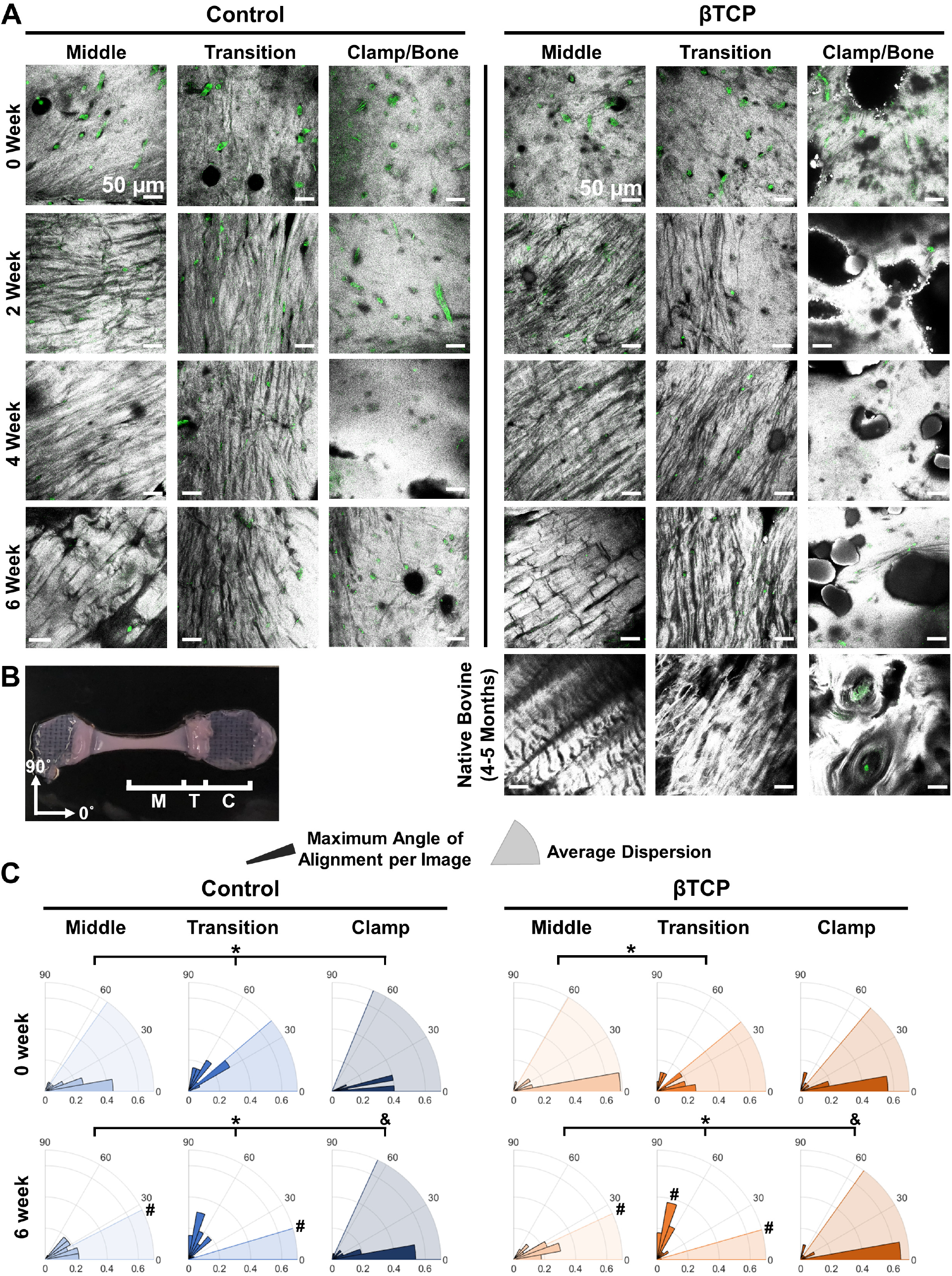
Clamping system drives zonal organization similar to immature bovine ligament-to-bone attachments (4-5 month). A) Confocal reflectance imaging reveals constructs are largely unorganized across all zones at 0 weeks, and by 6 weeks develop highly aligned parallel collagen fibers in the middle, smaller perpendicularly aligned fibers in the transition, and a dense, porous organization under the clamp. Grey = collagen, green = autofluorescence of cells; scale bar = 50 µm. B) representative image of construct at 6 weeks with middle (M), transition (T), and clamped (C) regions denoted and image analysis angle orientation. C) Image analysis of maximum angle of alignment of collagen fibers and degree of dispersion at 0 and 6 weeks. 3-5 images per zone of N = 4-11 constructs were analyzed for each timepoint. The maximum angle of alignment per image was binned at 10° increments and are represented as percentage of images per bin from 0° to 90°. The average dispersion of collagen fibers was determined by averaging all images per section and is represented as a single wedge from 0°. Image analysis confirmed constructs began largely unorganized at 0 week, characterized by fiber alignment close to 0° with large dispersion values. By 6 weeks, middle and transition zones were highly aligned in distinct directions with clamp zones remaining unorganized for both control and βTCP constructs. Significantly different *alignment angle, or ^&^fiber dispersion compared to bracket group, or ^#^compared to measurement at 0 week (p < 0.05).

#### 2.3.2 Scanning Electron Microscopy Analysis

SEM on 6 week constructs and native bovine ACL-to-bone was performed using a Hitachi SU-70 FE-SEM. Half-length sections were fixed in 10% formalin and stored in 70% ethanol. Prior to SEM analysis, ethanol content was serially increased to 100% and then constructs were dried via critical point drying. Constructs were mounted on 25 mm aluminum mounts with double sided carbon conductive tape and coated in platinum. Samples were imaged at a working distance of 15 mm, 5 kV of voltage, and 3000x magnification to observe collagen organization at the fibril level (<1 µm length scale).

#### 2.3.3 Polarized Picrosirius Red Imaging

Histological analysis was performed as previously described [41]. Briefly, fixed half-length sections and native bovine ACL-to-bone complexes were demineralized in 10% w/v ethylenediaminetetraacetic acid (EDTA) at pH of 7.4 and 4°C for 1 week, replacing EDTA and agitating daily. The constructs were then embedded into paraffin blocks, sectioned, and stained with picrosirius red. Constructs were imaged using a Nikon Eclipse Ts2R inverted microscope and Nikon Plan Fluor 10x/0.30 OFN25 Ph1 DLL objective under linear polarized light at 10x to observe collagen organization at the fascicle level (>100 µm length scale).

### 2.4 Tissue Composition

#### 2.4.1 Live/Dead Imaging

Live/Dead cell imaging was performed using LIVE/DEAD Viability/Cytotoxicity Kit ce(Invitrogen L3224). Half-samples (N = 3) were harvested and immediately incubated in a solution of 2 µM calcein AM and 4 µM ethidium homodimer-1 at room temperature for 30 minutes, washed twice with PBS, and imaged using a Zeiss 710 inverted LSM and LD C-Apochromat 40x/1.1 W Corr M27 objective. A 488 nm laser was split between confocal reflectance and fluorescence. Collagen was visualized as described in *2.3.1 Confocal Imaging Analysis*. Live cells were imaged at wavelengths of 499-557 nm. Dead cells were excited with a 561 nm laser, with images collected at 588-735 nm. Representative images were taken across the full length of each construct, with 2-3 images taken in each zone.

#### 2.4.2 Cell Morphology Analysis

Cell morphology analysis was performed in FIJI as previously described [55] on images taken during live/dead cell imaging. The live cell signal was separated from the collagen and dead cell signal for cell morphology analysis. The images were converted to black and white via the Threshold plugin, and area, shape descriptors, and centroid were measured. The Analyze Particles plugin was then applied to each image with a size filter of 50 to infinity µm^2^, circularity filter set from 0 to 1, and cells on the edge of the image were excluded. Circularity and aspect ratio (AR) data were recorded for each image as the average of all cells (∼10-20 cells) per image. Image data were pooled (2-3 images per section from N=3 constructs per time point) to determine average circularity and aspect ratio. Circularity is defined as 4π*area/perimeter^2^ with 1.0 indicating a perfect circle, and AR defined as major axis divided by minor axis fit to the centroid of each cell.

#### 2.4.3 Biochemical Analysis

Biochemical analyses for DNA, GAG, collagen content, and alkaline phosphatase activity were performed as previously described [41,44,51,52,56]. Briefly, tissue constructs were sectioned into middle, transition, and clamped zones upon removal from culture. For analysis of DNA, GAG, and collagen content, each respective section was weighed wet (WW), frozen, lyophilized, and weighed dry (DW) before being digested in 1.25 mg/ml papain solution at 60°C for 16 hours (N = 8-11). DNA, GAG, and collagen, were determined via Quantifluor dsDNA assay kit (Promega), modified 1,9-dimethylmethylene blue (DMMB) assay at pH 1.5 [57], and a modified hydroxyproline (hypro) assay [58]. Samples for Alkaline phosphatase (ALP) activity analysis were placed in 2% Triton-X immediately following dissection and frozen at -80°C (N = 5-8). To lyse cells, ALP samples were sonicated (Fisher Scientific, FB50) at a frequency of 20 kHz and amplitude of 60 for 10 seconds prior to testing. To remove residual βTCP, constructs were centrifuged at 1000 RPM at 4°C for 10 minutes, and the supernatant was used to analyze ALP activity via a modified ALP activity assay [56].

#### 2.4.4 Immunohistochemistry

Fluorescent immunohistochemistry (IHC) was performed to assess types I, II, and X collagen, and aggrecan localization. Engineered (N = 4) and native bovine ACL ligament-to-bone (N = 3) samples were fixed, demineralized as previously described in section *2.3.3 Polarized Picrosirius Red Imaging*, embedded in paraffin blocks and sectioned. Sections were treated with Proteinase K to retrieve antigens, blocked with 5% goat serum and incubated overnight with primary polyclonal rabbit antibodies in 5% goat serum at 1:150 dilution for types I (Abcam AB34710), II (Abcam AB34712), and X collagen (GeneTex GTX37732), and aggrecan (GeneTex GTX54920). Negative controls were incubated overnight in 5% goat serum. All sections were then incubated with goat anti-rabbit IgG secondary antibody labeled with Alexa Fluor 488 (Invitrogen A11008) at 1:200 dilution for 2 hours and cross labeled with DAPI at 1:1000 dilution. Stained sections were imaged using a Nikon Eclipse Ti2-E inverted microscope and Nikon Plan Fluor 10x/0.30 DIC L objective to observe protein localization.

### 2.5 Mechanical Analysis

Mechanical testing was performed as previously described [41,44,52]. Briefly, half-length tissue samples (N = 4-9) composed of the three zones were harvested and frozen to test for mechanical properties. All mechanical tests were performed using an EnduraTEC ElectroForce 3200 System and 250 g load cell. Constructs were thawed in PBS with EDTA-free protease inhibitor, measured, and secured in clamps with middle and clamped sections loaded into respective grips, ensuring the full transition and part of the middle and clamped sections were in between the grips. The constructs were loaded to failure at a rate of 0.75% strain/second, assuming quasi-static loading, and ensuring failure between grips. The linear region of the stress-strain curve was fit with a linear regression, ensuring r^2^ > 0.999, to determine the tensile modulus. The ultimate tensile strength (UTS) and strain at failure are the maximum stress point of the stress-strain curve.

### 2.6 Statistics

SPSS was used to confirm normality of data within each group and detect outliers using Shapiro-Wilk tests and Q-Q plots. After confirming normality, all image analyses, biochemical, and mechanical data were analyzed by 2 and 3-way ANOVA using Tukey’s *t*-test for *post hoc* analysis and *p* < 0.05 as the threshold for statistical significance. All data are represented as mean ± standard error (S.E.M.).

## 3. Results

### 3.1 Gross Tissue Morphology

Gross level inspection revealed both control and βTCP treated constructs had similar contraction over 6 weeks of culture (**Figure 2B**). Specifically, constructs maintained their size and shape through ∼4 weeks, followed by significant contraction in the mid-section by 6 weeks.

### 3.2 Zonal and Hierarchical Collagen Organization

#### 3.2.1 Zonal Collagen Fiber Organization

Confocal reflectance of both control and βTCP constructs revealed development of 3 unique zones of collagen organization in the middle, transition, and clamped regions, which had similar organization to immature (4-5 months old) native bovine tissue in the ligament, transition, and bone, respectively (**Figure 3A**). Specifically, at 0 weeks both control and βTCP constructs consisted of largely unorganized collagen across all three zones. By 2 weeks, distinct zonal organizations began to form, with highly aligned parallel collagen fibers in the mid-section, perpendicular aligned fibers in the transition, and a denser unorganized, porous network in the clamped region. By 6 weeks, this organization had further matured, with larger collagen fiber formation in the middle, similar to previous work which evaluated fiber formation in the mid-section alone [41]. Further, by 6 weeks βTCP constructs displayed increased porosity under the clamp and more robust fiber formation in the transition compared to control samples (**Figure 3A**).

Image analysis of collagen fiber organization confirmed that at 0 weeks, control and βTCP constructs had similar organization between middle and clamped regions with maximum angles of alignment between 10-20° and large values of dispersions (∼50-60°). Interestingly, after 24 hours of culture (0 week timepoint), the transition region for control and βTCP constructs had a higher mean fiber alignment compared to middle and clamped regions, with alignments of 49.3 ± 4.3° in control and 35.3 ± 8.4° in βTCP transitions, suggesting early re-organization (**Figure 3B** and **Supplemental Figure 2**). By 6 weeks, the middle region of control and βTCP constructs had maximum angles of alignment at 24.3 ± 2.8° and 23.2 ± 2.8°, respectively, while the transition region of control and βTCP constructs had a significantly different fiber alignment, with fibers oriented at 65.0 ± 3.0° and 66.9 ± 3.5°, respectively. Further, both control and βTCP constructs by 6 weeks had significantly less dispersion (∼20-30° dispersion) in the middle and transition sections when compared to 0 week (**Figure 3B** and **Supplemental Figure 2**). In contrast, the clamped region for both control and βTCP constructs maintained a steady alignment, with average alignments of 17.9 ± 4.0° for controls and 19.2° ± 5.1° for βTCP constructs by 6 weeks. Additionally, the clamped region for both control and βTCP constructs maintained large degrees of dispersion, with dispersion averages of 66.2° ± 7.4° for control constructs and 56.2° ± 5.2° for βTCP constructs by 6 weeks (**Figure 3B** and **Supplemental Figure 2**). Collectively, this data demonstrates both control and βTCP constructs develop more organized fibers, aligned at ∼25° in the middle region, more organized fibers, aligned at ∼65° in the transition, and largely unorganized, un-aligned collagen matrix in the clamped section (**Figure 3B**).

#### 3.2.3 Hierarchical Organization

Similar to fiber level organization evaluated by confocal (1-100 µm length scale), SEM and picrosirius red staining revealed both control and βTCP constructs maintained native-like zonal organization at the fibril (<1 µm length scale) and fascicle level (>100 µm length scale), respectively (**Figure 4**). At 6 weeks, SEM analysis revealed fibril level organization similar to confocal analysis, with highly aligned parallel collagen fibrils in the mid-section, perpendicularly aligned fibrils in the transition, and unorganized, dense matrix in the clamped region, similar to native immature tissue. Additionally, βTCP samples displayed sheet-like mineralization [59] under the clamp similar to native immature bovine tissue (**Figure 4A**). Picrosirius red staining, performed at lower magnification than confocal, revealed both control and βTCP constructs maintained aligned parallel collagen fibers in the mid-section and perpendicularly aligned collagen fibers in the transition, similar to native immature tissue; however engineered constructs largely lacked the larger collagen fascicles observed in native tissue. Under the clamp, both control and βTCP constructs had collagen fibers aligned around pores, similar to native bone (**Figure 4B**).

**Figure 4:**
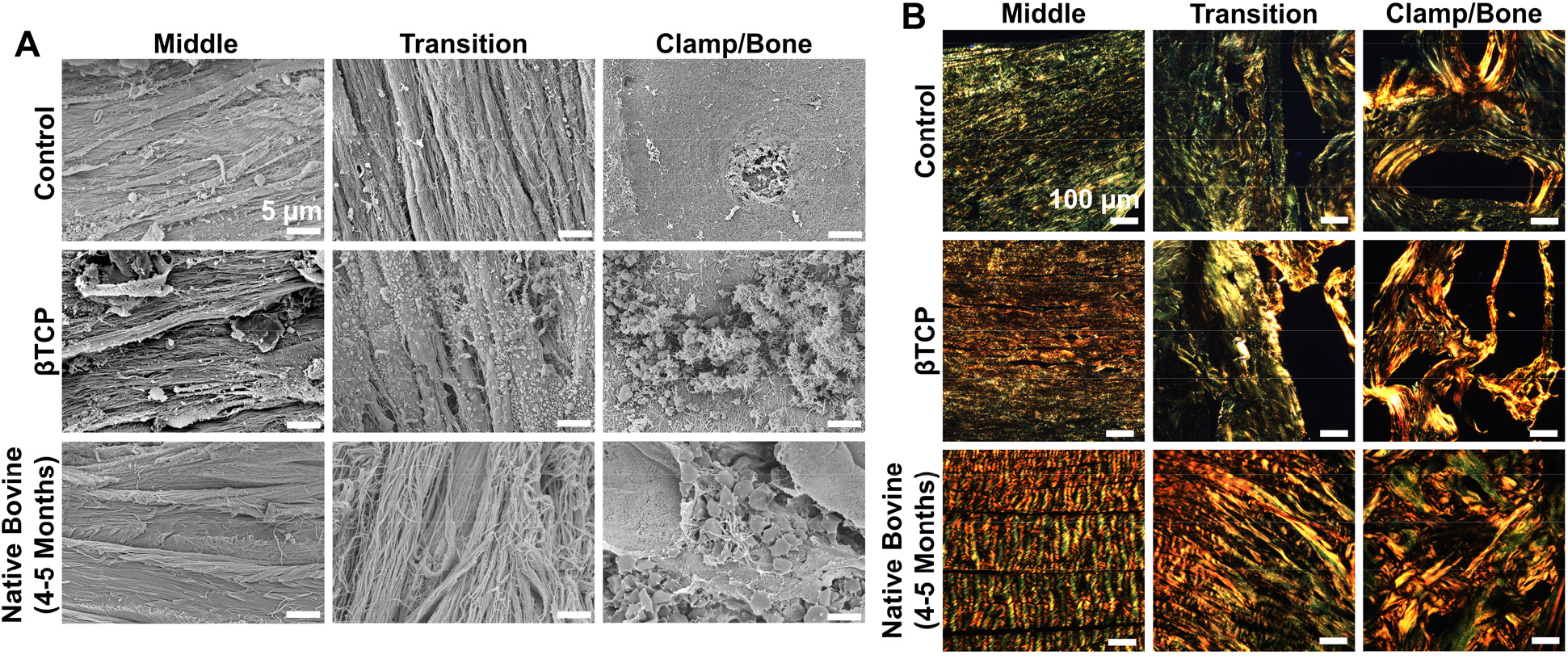
Six week constructs maintain zonal organization similar to immature bovine ligament-to-bone attachments at the fibril and fascicle length-scale. A) Fibril length-scale organization assessed via SEM (scale bar = 5 µm) and B) fascicle length-scale organization evaluated by picrosirius red staining imaged with polarized light (scale bar = 100 µm). Middle and transition zones maintained parallel and perpendicular fiber alignment at both magnifications, while clamped regions revealed A) sheet-like mineralization in βTCP constructs and B) collagen aligned around pores in both sets of constructs.

### 3.3 Zonal Tissue Composition

#### 3.3.1 Zonal Cellular Distribution

Both control and βTCP constructs developed similar zonal DNA concentrations by 2 weeks of culture. Specifically, both sets of constructs had a significant increase in DNA by 2 weeks in the middle and transition zones, while the clamped zone remained unchanged. This zonal distribution of DNA remained constant throughout the rest of culture with the exception of the βTCP clamped section which significantly increased at 4 weeks and subsequently returned to baseline by 6 weeks (**Figure 5A**). Live/Dead staining revealed cells remained viable in all zones throughout culture, with no noticeable decrease in percent viability under the clamp (**Supplemental Figure 3A**). Further, cell morphology analysis of live cells revealed that cells in both sets of constructs became more elongated in the middle and transition zone, suggesting a more ligament fibroblast phenotype, and more rounded under the clamp, suggesting a more chondrocyte-like phenotype (**Supplemental Figure 3B**). Specifically, by 6 weeks, control constructs had significantly different cellular aspect ratios in each zone, with the most elongated (higher aspect ratio) cells in the transition. Similarly, βTCP constructs at 6 weeks had significantly more elongated cells in the transition zone compared to the clamped zone. Further, control constructs maintained significantly more circular cells in the clamp region from 2 weeks on, while βTCP constructs had more circular cells in the clamped region starting at 4 weeks. Interestingly, by 6 weeks cells in the transition zone of βTCP constructs had similar circularity as cells in the clamped region (**Supplemental Figure 3B**).

**Figure 5:**
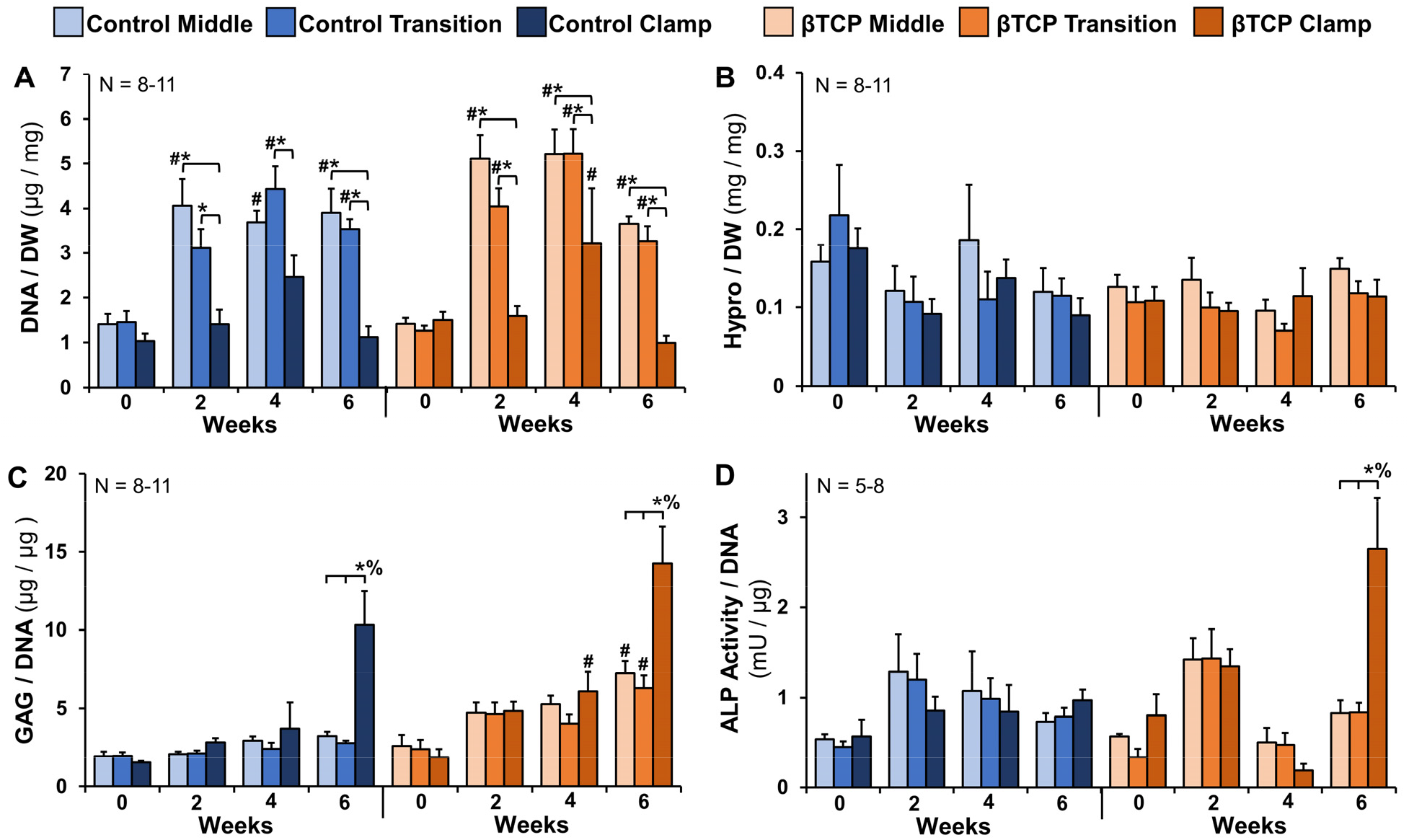
Compositional measurements of the middle, transition, and clamped zones for control and βTCP constructs. A) DNA normalized to dry weight (DW); B) hydroxyproline (hypro), as a measure for collagen content, normalized to dry weight; C) Glycosaminoglycans (GAGs) normalized to DNA; and D) ALP activity measured as nmol p-nitrophenol substrate catalyzed per minute (mU) normalized to DNA. DNA and collagen content remained largely constant throughout culture, while GAG and ALP activity significantly increased in clamped regions at 6 weeks. Significance compared to *bracket, ^#^0 week, or ^%^respective zone at all timepoints (p < 0.05). Data shown as mean ± S.E.M.

#### 3.3.2 Zonal Biochemical Distribution

Since constructs were composed of high-density collagen gels, collagen content, represented by hydroxyproline (hypro), is reported normalized to dry weight, while GAG and ALP, which are only added to the system by cells, are reported normalized to DNA. Collagen content for both control and βTCP constructs remained relatively constant throughout culture for all sections (**Figure 5B**). GAG content, a marker of more chondrogenic tissue, remained relatively constant in both sets of constructs until 6 weeks. At 6 weeks, both control and βTCP constructs had a significant 2-3 fold increase in GAG/DNA in the clamped region compared to mid and transition zones. Interestingly, βTCP constructs accumulated more GAG/DNA than controls, with all three zones of βTCP constructs having increased GAG accumulation at 6 weeks over respective 0 week zones (**Figure 5C**). ALP activity, a marker of more osteogenic cells, remained relatively constant throughout culture for both control and βTCP constructs. However, at 6 weeks βTCP constructs significantly increased ALP activity in the clamped region ∼2.5 fold over all other zones of control and βTCP samples (**Figure 5D**).

Six week constructs were evaluated for spatial distribution of type I, II, and X collagen, and aggrecan (**Figure 6**). As expected, type I collagen was dispersed uniformly throughout the middle, transition, and clamped zones for both sets of constructs (**Figure 6A**). Type II and X collagen were primarily localized to the clamped region for both control and βTCP constructs, with spotty accumulation of type II collagen in the transition (**Figure 6B-C**). Aggrecan was largely isolated to the clamped sections for both control and βTCP constructs, with reduced spotty accumulation in the transition as well (**Figure 6D**). While type II collagen, type X collagen, and aggrecan distribution do not match the immature (4-5 month old) bovine ligament-to-bone tissue, this distribution does match previously reported neonatal bovine distribution (1-7 days [36]).

**Figure 6:**
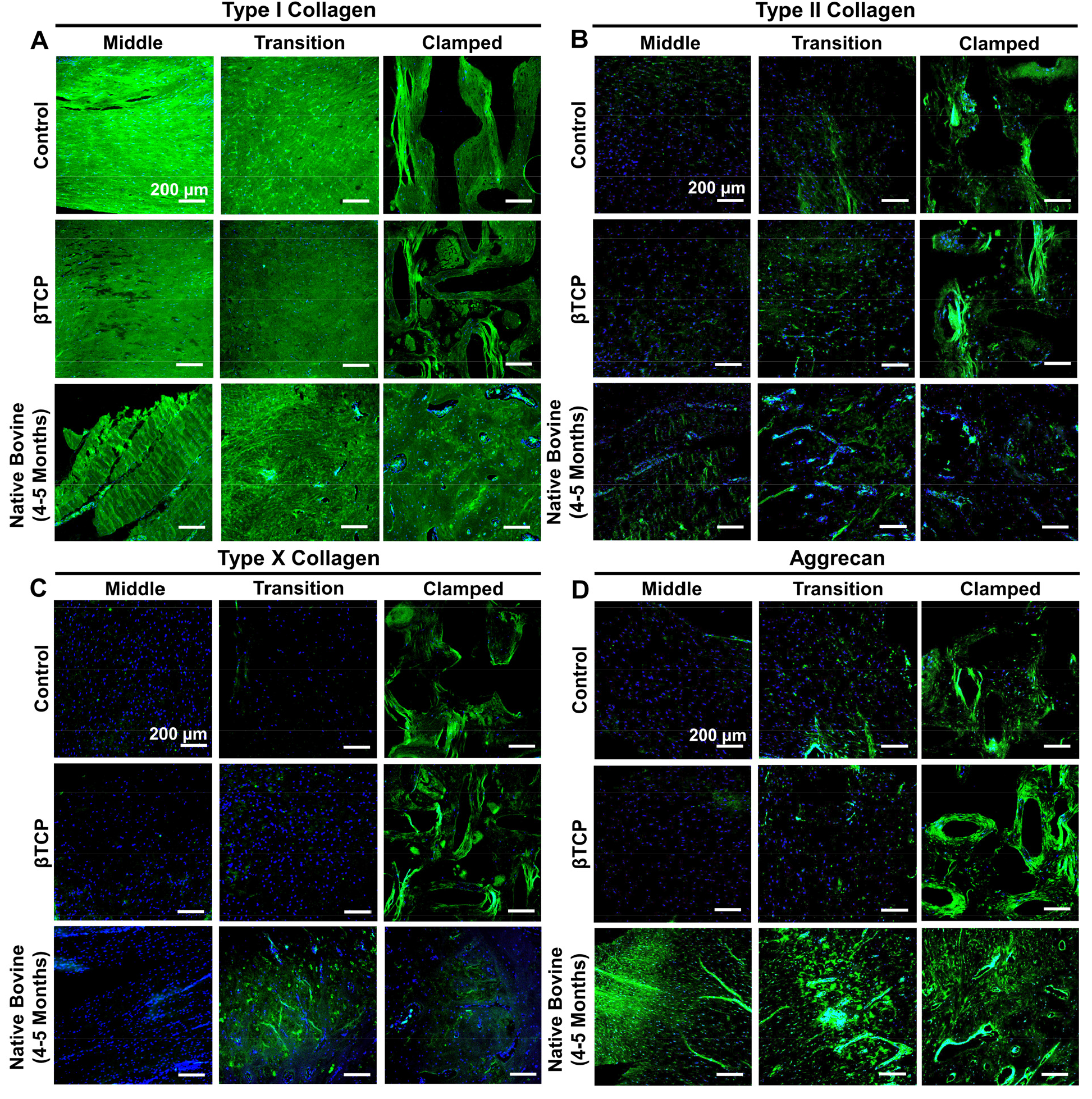
Zonal immunohistochemistry of control and βTCP constructs at 6 weeks compared to native immature bovine ligament-to-bone tissue. A) Type I collagen; B) Type II collagen; C) Type X collagen; and D) Aggrecan. FITC =respective protein, DAPI = cells, scale bars = 200 µm.

### 3.4 Mechanical Properties

Tensile modulus across the transition zone of βTCP constructs improved significantly from 0 to 6 weeks, reaching ∼2.5 MPa by the end of culture, whereas the tensile modulus of control constructs reached only ∼1.5 MPa by 6 weeks, significantly less compared to βTCP constructs (**Figure 7B**). By 6 weeks, the transition tensile moduli of control and βTCP constructs surpassed reported bovine neonatal ACL entheses moduli (∼1 MPa [4]) and matched properties of immature bovine ACL (∼1-3 MPa [60]). By 6 weeks, control and βTCP constructs had similar ultimate tensile strength (UTS, **Figure 7C**), however, βTCP constructs had significantly lower ultimate strain properties compared to control constructs, suggesting further maturation (**Figure 7D**).

**Figure 7:**
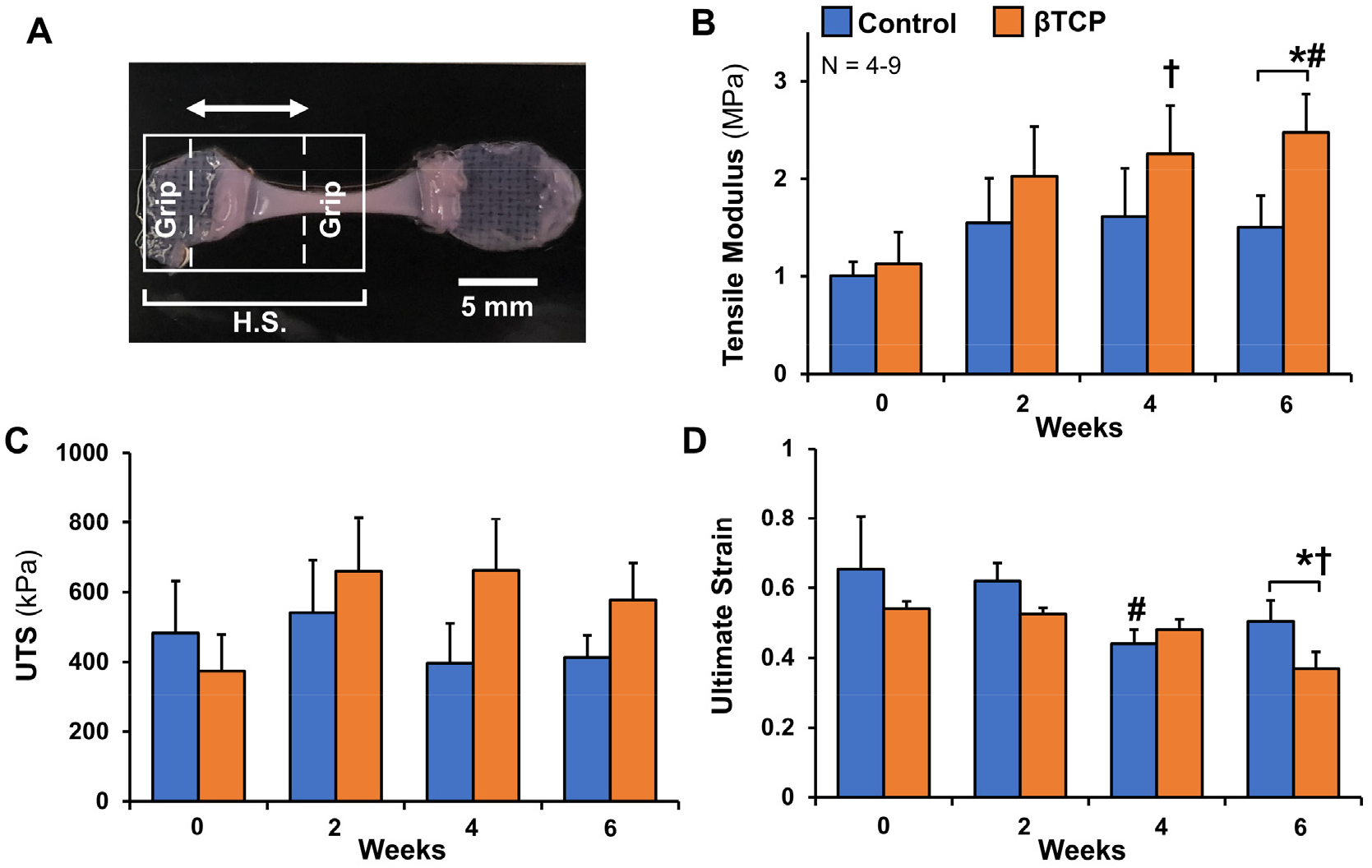
Mechanical testing revealed βTCP constructs developed increased tensile modulus and decreased strain at failure across the transition compared to control by 6 weeks, suggesting increased tissue maturation. A) Depiction of half-length samples (H.S.) loaded into the grips for tensile testing across the transition zone (arrow). B) Tensile modulus, C) ultimate tensile strength (UTS), and D) ultimate tensile strain across the construct transition zone. Significance compared to *bracket or ^#^0 week (p < 0.05). Trending compared to ^†^0 week (p < 0.1). Data shown as mean ± S.E.M.

## 4. Discussion

Fibrocartilaginous entheses are structurally complex tissues 100 µm to 1 mm wide that translate load from elastic ligaments to stiff bone due to an intricate zonal organization with gradients in collagen fiber organization, matrix composition, and cell phenotype [1–4]. Currently, these gradients, necessary for long-term mechanical function, are not recreated in soft tissue-to-bone healing or engineered replacements, leading to high failure rates and a lack of repair options [2–4]. Research has primarily focused on top-down scaffold design using aligned synthetic or natural fibers and induced spatial mineralization [3,21,22,25,61,62]. These techniques attempt to replicate mature ligament and bone organization; however, they may actually impede development of the physiological gradient of the enthesis since aligned fibers and mineralization are not present until later in development. Here we sought to use a bottom-up approach, inspired by development, to generate the complex gradients of the enthesis. This study demonstrates harnessing cellular contraction forces with compressive boundary conditions produces a tensile-compressive interface that results in 3 unique zones of collagen organization and zonal matrix composition similar to immature bovine ACL-to-bone attachments. Further, the addition of βTCP under the clamp, a known osteoconductive agent and aid in mineralization [46,47], enhances the zonal organization, composition, and mechanical properties of the engineered entheses.

The developing enthesis is characterized by highly aligned ligament-like tissue transitioning into cartilage, which later matures into fibrocartilage and bone (**Figure 1**, [32,35,36]). Previously, we demonstrated that the external clamps in our culture system restrict cellular contraction length wise, producing a tensile load cells pull against in the mid-section. Ultimately, this harnessing of cellular contraction guides neonatal ligament fibroblasts to form aligned, native sized ACL collagen fibers 30 µm in diameter, which group together to form ∼200 µm diameter fiber bundles by 6 weeks of culture [41]. In this study, we further analyzed this system to demonstrate that the compressive environment under the clamps produces tissue with a more unorganized collagen matrix, increased GAG accumulation, and localized type II and type X collagen accumulation, similar to the cartilage-like tissue that composes the early neonatal enthesis [36]. As discussed in the introduction, traditionally, tensile loads are often fibrogenic, increasing type I collagen synthesis and tensile properties, while compressive loads are chondrogenic, increasing GAG and type II collagen synthesis. Our findings mirror this in the development of the middle and clamped sections, however the more unexpected changes occurred in the transition interface between the tensile middle and compressive clamps.

Collagen fibers in the transition section aligned perpendicular to the middle section with time in culture, ultimately producing three unique zones of organization. Specifically, by 6 weeks for both βTCP and control constructs, collagen in the middle section aligned at 20°-30°, collagen in the transition aligned at a significantly different angle of ∼65°, and collagen under the clamp remained largely unorganized with significantly higher degrees of dispersion. This organization was maintained across hierarchical levels with similar organization seen at both fibril and fiber length scales (**Figures 3A and 4A**). This zonal organization mirrors early ligament-to-bone development where the neonatal enthesis is largely composed of homogenously unorganized type I collagen fibrils [28] which then align perpendicular to the ligament proper during postnatal development (**Figure 1**) [32]. As noted in the introduction, as the enthesis further matures, collagen shifts from being perpendicular to the ligament to a more diffuse orientation parallel with the ligament [25,35,36]. This final collagen organization is thought to be critical to reducing stress concentrations and providing functional translation of load [30,31,33,34,37].

Collagen organization under the clamp remained largely unorganized; however, with time in culture there was a noticeable increase in larger pores (**Figure 3A**) with collagen oriented circumferentially around pores (**Figure 4B**), suggesting a transition from more cartilage like tissue to early bone formation [63]. Further, SEM (**Figure 4A**) revealed βTCP constructs had sheet-like mineralization of the collagen fibrils under the clamp similar to native tissue [59]. The combination of porosity and mineralization observed under the clamp for βTCP constructs is consistent with the expected subchondral bone structure consisting of type I collagen infiltrated with calcium phosphate mineral arranged in a porous network [1,64].

In addition to driving zonal collagen organization, cell shape changes were observed across the three zones with time in culture as well. At 0 weeks, cells largely maintained similar circularity across all zones. By 6 weeks, control constructs revealed more elongated cells similar to ligament fibroblasts in the middle and transition zones, and more circular cells under the clamps similar to chondrocytes (**Supplemental Figure 3B**). As mentioned previously, the enthesis is characterized by a wide array of cell types. Early in development the enthesis is primarily composed of ligament fibroblasts and chondrocytes, which later transition to include fibrochondrocytes, hypertrophic chondrocytes, osteoblasts, osteoclasts, and osteocytes. Interestingly, with the addition of βTCP under the clamp, cells in the transition zone demonstrated increased circularity compared to control constructs, possibly suggesting a shift to a more fibrocartilage or cartilage like phenotype [44,55]. Of note, in this study we use immature cells isolated from the entire ACL of 0-5 months old bovine, thus it is possible our cell population either begins with these different cell types present or begins with a sub-population of progenitor cells that supports the observed changes in cell type. While cell aspect ratio is a known measure of cell morphology and phenotype [55], further analysis should be performed to verify cellular changes.

Mirroring observed cell shape changes, matrix composition changed as well. As mentioned, the accumulation of GAG increased under the clamp at 6 weeks for both control and βTCP constructs (**Figure 5C**), suggesting the development of a more cartilage-like tissue. These results mimic GAG production in early development of native entheses, which has been shown to be influenced by compressive loads [42,43]. More specifically, the bone begins as cartilage with high levels of GAG [36] and with further development shifts to the interface region due to complex forces generated at the ligament-to-bone interface [32,35]. The compressive environment induced by the clamp may drive the increase in GAG and cartilage-like tissue; however, it is possible there is a hypoxic environment under the clamp as well, which is also known to induce cartilage-like tissue [65,66]. However, in our culture system there is a stainless steel mesh insert under the clamp which should reduce hypoxia and aid nutrient diffusion, thus we believe the development of cartilage-like tissue is primarily due to the compressive environment of the clamp.

The addition of βTCP under the clamp resulted in increased GAG accumulation under the clamp compared to control constructs and increased ALP activity by 6 weeks as well (**Figure 5D**). ALP is a well characterized early marker in bone development and is implicated in the development of the mineralized tissues in the enthesis [35]. It has been suggested that ALP expression prior to the initialization of osteoblast mineralization is involved in the preparation of ECM for proper mineral deposition [67]. Previous studies have demonstrated that increased ALP activity is inversely related to proliferation and that as proliferation decreases, or is halted, ALP activity significantly increases [68–70]. This relationship can be seen in the proliferation under the clamp at 4 weeks followed by the subsequent decrease in DNA at 6 weeks (**Figure 5A**). The increases in GAG accumulation and ALP activity following the decrease in DNA from 4 to 6 weeks in the clamped region suggest tissue maturation via the early stages of endochondral ossification, mirroring early native enthesis development.

In addition to the changes in cell shape, GAG accumulation, and ALP activity, there was also a shift in the types of collagen accumulated under the clamp. While the general amount of collagen remained consistent across all zones with time in culture (**Figure 5B**), differences in types of collagen were found within the zones. Notably, type II and type X collagen were largely localized under the clamp for both control and βTCP constructs at 6 weeks. Additionally, aggrecan was also localized under the clamp for both sets of constructs at 6 weeks, with spotty accumulation observed in the transition sections (**Figure 6D**). In early enthesis development, the cartilage/bone region is characterized by type II collagen and aggrecan, with type X collagen accumulating via endochondral ossification [36]. With further maturation, type II collagen, type X collagen, and aggrecan shift from the cartilage/bone region to the developing fibrocartilage interface, ultimately resulting in gradients forming unmineralized and mineralized fibrocartilage zones [1,3,4,33]. Interestingly, the transition displayed some aggrecan and type II collagen accumulation but was largely void of type X collagen, suggesting the need for further maturation. However, the accumulation of type II collagen, type X collagen, and aggrecan under the clamp, in addition to increased GAG accumulation and ALP activity, further suggests that clamping initiates early stages of endochondral ossification mirroring native development.

The presence of type II and type X collagen under the clamp in βTCP constructs suggests possible maturation to fibrillar mineralization, similar to the sheet-like mineralization observed under the clamp in SEM analysis (**Figure 4A**). In this study, βTCP was chosen for investigation as it has been demonstrated to release a greater amount of Ca^2+^ ions *in vitro* than hydroxyapatite [48]. The breakdown of βTCP combined with the increase in ALP activity and GAG accumulation under the clamp likely plays a role in the sheet-like mineralization observed in SEM, suggesting the possible initiation of the mineralization phase in βTCP samples. The process of mineralization during endochondral ossification has been shown to be influenced by the presence and accumulation of proteoglycans, specifically GAGs [71,72]. The mineralization phase begins when hypertrophic chondrocytes and osteoblasts release matrix vesicles (MV) containing Ca^2+^ ions and proteins such as phosphatases [71–74]. The MVs bind with accumulated proteoglycans, and due to the negative charge of proteoglycans are able to immobilize the Ca^2+^ ions stored within the MVs [71]. Perivesicular and intravesicular phosphate ion (PO_4_^3-^) concentration increases with alkaline phosphatase activity [73], and then these phosphate ions cross the vesicular membrane to trigger nucleation and formation of hydroxyapatite crystals [71,73]. Further, type II and type X collagen are believed to localize and bind to the outer surface of the MVs and may serve as a bridge for mineral propagation and collagen fibril mineralization [73].

With the zonal changes in matrix organization and composition, the mechanical properties of the engineered entheses changed with time in culture as well. The addition of βTCP under the clamp resulted in a significant increase in the tensile modulus of the enthesis transition zone by 6 weeks compared to 0 week and 6 week control constructs. Ultimately, the tensile modulus of the βTCP constructs reach ∼2.5 MPa, surpassing the moduli reported for neonatal bovine entheses (0-1 week old [4]) and matching reported moduli of neonatal bovine ACL (1 week old [60]). Further, βTCP constructs had a significant decrease in ultimate strain by 6 weeks, indicating further matrix maturation. The majority of ACL injuries are reported to occur at the enthesis due to the complex loading environment [3,4], and the majority of graft repairs fail at the enthesis due to a lack of regeneration [3], demonstrating the need for strong attachments. Previously, we have reported that the tensile modulus of the middle section of our constructs reaches ∼1 MPa by 6 weeks of culture [41]. Both the control and βTCP constructs develop transitions with tensile moduli greater than 1 MPa by 6 weeks of culture, suggesting the development of a stronger attachment and lack of structural weak point at the interface. Interestingly, the tensile modulus for control constructs plateaued at 2 weeks only reaching ∼1.5 MPa despite observed organizational changes occurring between 2 and 6 weeks, suggesting a possible disconnect in structure and mechanics. Changes in matrix composition, including GAG accumulation and ALP activity, occur primarily at 6 weeks suggesting that tissue maturation is ongoing at the end of culture and that more time in culture may be needed to further enhance mechanical properties.

This study demonstrated that compressive boundary clamps that restrict cellular contraction across the construct, producing a zonal tensile-compressive environment that guides cells to produce 3 unique zones of collagen organization, and zonal GAG, type II collagen, and type X collagen accumulation, ultimately resulting in engineered entheses with similar organization and composition as immature bovine ACL-to-bone tissue. Further, the addition of βTCP under the clamp enhanced the development and maturation of these entheses leading to increased GAG production and sheet-like mineralization under the clamp, and improved mechanical properties across the transition zone, suggesting the initiation of endochondral ossification and the development of more robust attachments. While zonal collagen organization at the fibril and fiber level (<100 µm length scale) aligned closely with native immature tissue, higher-level fascicle organization (>100 µm length scale) was lacking, suggesting the need for further maturation. Additionally, while zonal accumulation of type II collagen, type X collagen, and aggrecan under the clamp match expected zonal accumulation for early postnatal entheses [1–4], further maturation is needed to shift these proteins into the transition zone, as is observed in late postnatal entheses [1,3,4,33]. Collectively, this culture system uses a bottom-up approach to produce tissue which closely mirrors neonatal to early postnatal enthesis development. However, improvements are needed to further mature the tissues. Biochemical and mechanical stimulation are well established to further improve engineered tissue development [21,43–45,61,62,75,76]. The addition of conditioned media or tissue specific growth factors has been shown to enhance zonal soft tissue-to-bone development [62,75,76]. Further, mechanical cues are well established to be essential to enthesis development [29,38,39], and thus addition of mechanical load could also further drive maturation of these tissues. Despite these limitations, the entheses developed in this study are some of the most organized ligament-to-bone entheses developed to date. This culture system, which closely mimics enthesis development, holds great promise for better understanding the cues which drive enthesis formation *in vivo* and to shed light on how to better drive enthesis regeneration following graft repair.

## 5. Conclusions

This study provides new insight into how βTCP and cell-mediated tensile-compressive boundary conditions synergistically drive zonal gradients in organization, mineralization, and matrix composition, ultimately producing some of the most organized engineered enthesis constructs to date. These constructs demonstrate great promise as functional ACL replacements with zonal organization, composition, and mechanics similar to immature bovine ACL-to-bone complexes. Further, the *in vitro* system used to develop these constructs, which mirrors enthesis development, is also a promising tool to further investigate the effects of mechanical loading, chemical signaling, and cellular differentiation in enthesis maturation.

## Acknowledgements

The authors would like to thank Dr. Rene Olivares-Navarrete, Dr. Hank Donahue, Dr. Carl Sayer, Dr. Dmitry Pestov, Dr. Joseph Turner, and O.J. Juhl for their assistance with this study. The authors acknowledge the use of facilities within the Nanomaterials Characterization Core and Chemical Research Instrumentation Facility at Virginia Commonwealth University. The authors acknowledge that services and products in support of the research project were generated by the Virginia Commonwealth University Cancer Mouse Models Core Laboratory, supported, in part, with funding from NIH-NCI Cancer Center Support Grant P30 CA016059. Finally, this work was supported, in part, by a grant from the Orthoregeneration Network (ON) Foundation, Switzerland (19-037) and PI Start-up Funds.

