## Supplemental Figures for "Driving Native-like Zonal Enthesis Formation in Engineered Ligaments Using Mechanical Boundary Conditions and β-Tricalcium Phosphate"

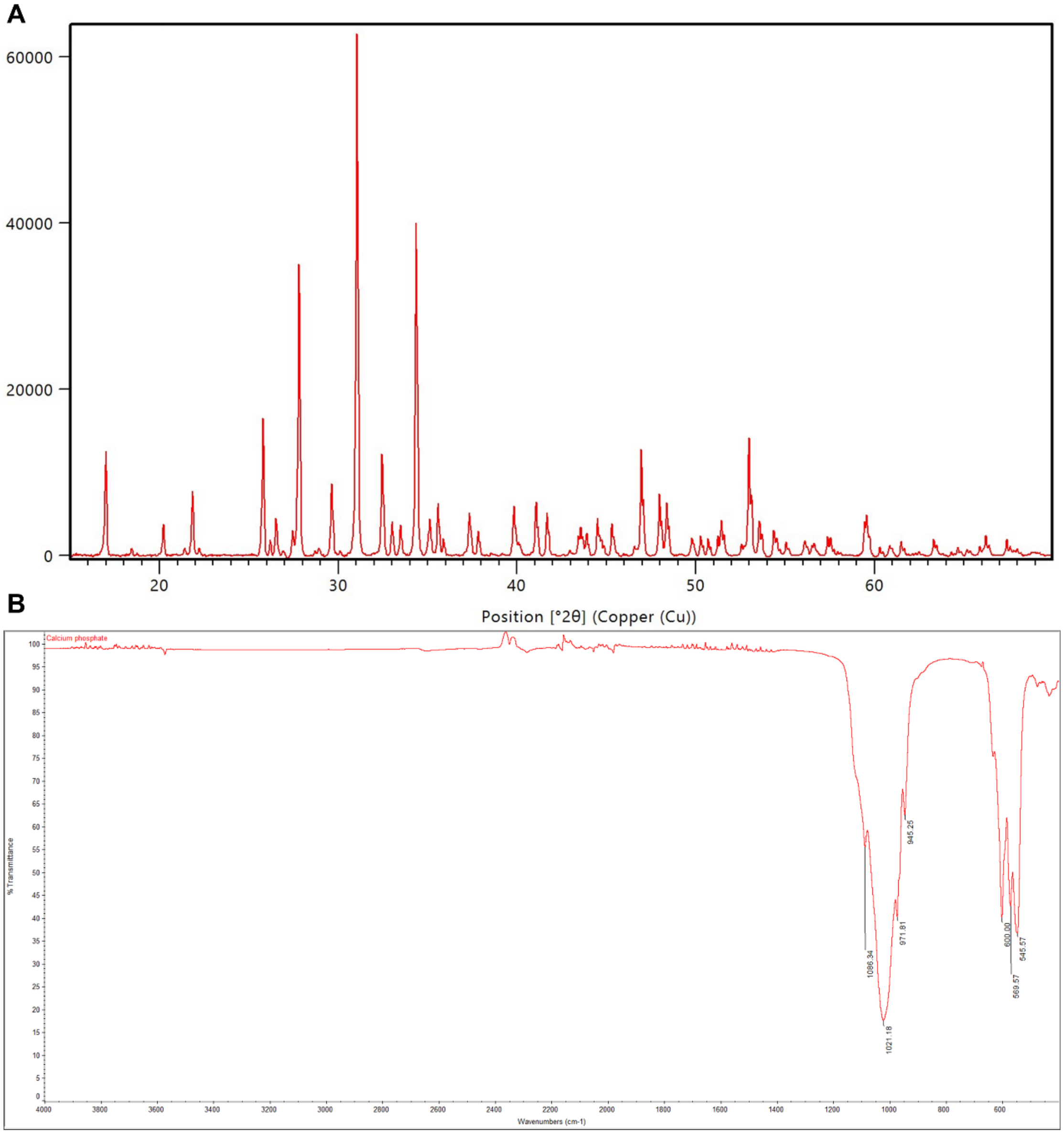


**Supplemental Figure 1:** A) XRD and B) FTIR of synthesized βTCP display characteristic peaks and absorption bands of βTCP.

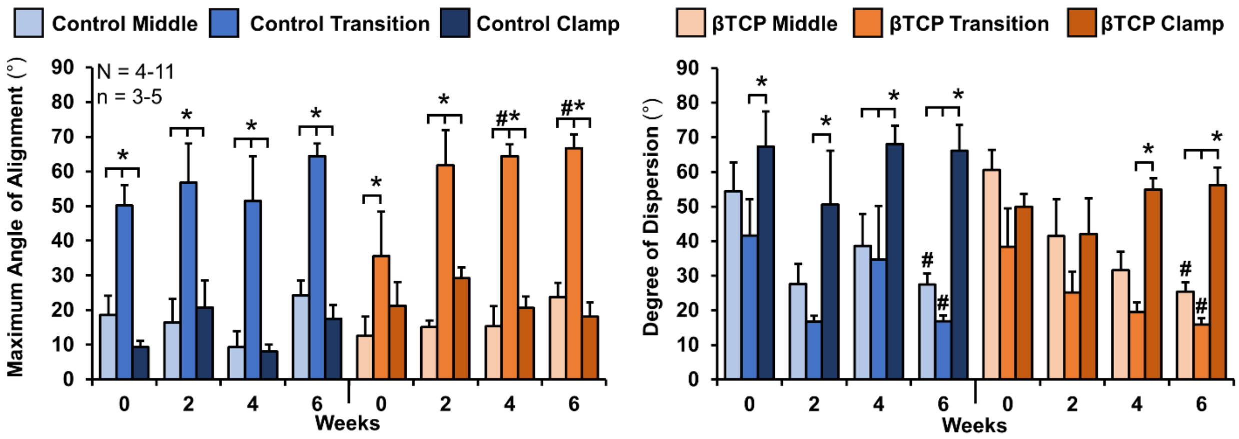


**Supplemental Figure 2:** Confocal image analysis of the maximum angle of collagen fiber alignment and degree of dispersion of collagen fibers for 0 through 6 weeks. Collectively, both control and βTCP constructs develop organized fibers aligned at ~25° in the middle region and organized fibers aligned at ~65° in the transition, with significantly reduced dispersion by 6 weeks. In the clamped section, both sets of constructs maintained largely unorganized, un-aligned collagen matrix, with significantly larger degrees of dispersion compared to middle and transition zones by 6 weeks. 3-5 images per zone for each construct were analyzed, with 4-11 constructs analyzed for each time point. Significance compared to *bracket or #0 week ($p$<0.05). Data shown as mean ± S.E.M.

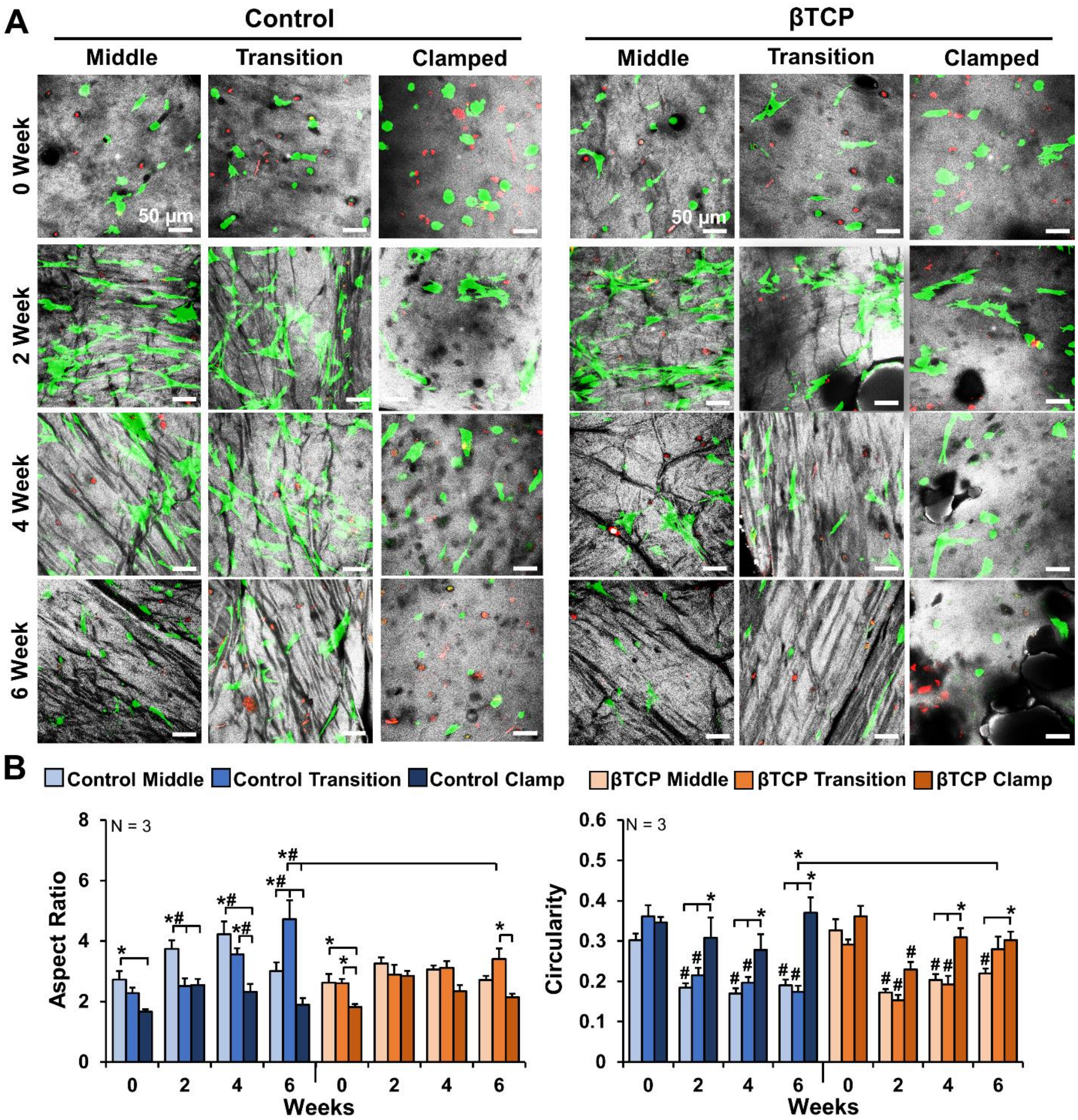


**Supplemental Figure 3**: A) Live/dead cell analysis reveals cells maintain similar viability across all zones throughout the 6 weeks and undergo a morphological shifts in cell shape between the zones. Grey is collagen, green is live cells, and red is dead cells; scale bar is 50 µm. B) Image analysis of live cells revealed cells in the middle and transition zone became more elongated with higher aspect ratios, while cells in the clamped region were significantly more circular by 6 weeks. Average measurements of 10-20 cells per image, from 2-3 images per zone of N = 3 constructs were analyzed for each time point. Significance compared to *bracket or #0 week ($p < 0.05$).
